# Homology-directed repair involves multiple strand invasion cycles in fission yeast

**DOI:** 10.1101/2020.05.03.074468

**Authors:** Amanda J. Vines, Kenneth Cox, Bryan A. Leland, Megan C. King

## Abstract

Homology-directed repair of DNA double-strand breaks (DSBs) can be a highly faithful pathway. Non-crossover repair dominates in mitotically growing cells, likely through a preference for synthesis-dependent strand annealing (SDSA). While genetic studies highlight a key role for the RecQ helicase BLM/Rqh1 (in human and *S. pombe* cells, respectively) in promoting noncrossover repair, how homology-directed repair mechanism choice is orchestrated in time and space is not well understood. Here, we develop a microscopy-based assay in living fission yeast to determine the dynamics and kinetics of an engineered, site-specific interhomologue repair event. We observe highly efficient homology search and homology-directed repair in this system. Surprisingly, we find that the initial distance between the DSB and the donor sequence does not correlate with the duration of repair. Instead, we observe that repair is likely to involve multiple site-specific and Rad51-dependent co-localization events between the DSB and donor sequence, suggesting that efficient interhomologue repair in fission yeast often involves multiple strand invasion events. By contrast, we find that loss of Rqh1 leads to successful repair through a single strand invasion event, suggesting that multiple strand invasion cycles reflect ongoing SDSA. However, failure to repair is also more likely in *rqh1Δ* cells, which could reflect increased strand invasion at non-homologous sites. This work has implications for the molecular etiology of Bloom syndrome, caused by mutations in BLM and characterized by aberrant sister chromatid crossovers and inefficient repair.

## Introduction

Homology-directed repair (HDR) is a conserved, high-fidelity mechanism for repairing DNA double strand breaks (DSBs). Following recognition of the DSB by the MRN complex and 5’ to 3’ exonuclease-dependent end resection, faithful repair by HR requires a Rad51-dependent homology search by the resultant nucleoprotein filament to locate a homologous donor sequence as a template. The sister chromatid available after replication is the most common template for repair (San Filippo *et al*., 2008; Mimitou and Symington, 2009). In this case, the template is identical to the original sequence and homology search is likely to be temporally and spatially efficient due to sister chromatid cohesion(Seeber *et al*., 2016; Haber, 2018). The homologous chromosome or an ectopic sequence can also be used as templates for repair, involving an expected less efficient homology search (Pâques and Haber, 1999; Mehta and Haber, 2014), but these can lead to loss of heterozygosity or genome instability(Renkawitz *et al*., 2014). Therefore, high fidelity repair hinges on the accurate choice of a homologous donor.

A successful long-range homology search (i.e. with a non-sister chromatid donor) requires that (1) the distant DSB and donor loci are able to encounter one another within the nucleus; (2) the Rad51-bound nucleoprotein filament can drive strand invasion of potential donors (leading to formation of a displacement (D-) loop; and (3) the homologous sequence is used as the template for new synthesis. Chromatin mobility likely facilitates the encounter rate and often increases upon DSB induction both locally at the DSB and globally(Miné-Hattab and Rothstein, 2012; Seeber *et al*., 2013). The degree of induced mobility may be influenced by the type of damage induction (irradiation, DNA damaging drugs such as zeocin or site-specific nuclease induction), cell ploidy (chromatin density), the number of DSBs and whether a DSB has persisted long enough to activate checkpoint arrest(Miné-Hattab *et al*., 2017; Zimmer and Fabre, 2019). Indeed, chromatin mobility can be induced by activation of the checkpoint response in the absence of damage (Bonilla *et al*., 2008). Notably, an initial decrease in mobility within the first hour following DSB induction has been observed in budding yeast with a single DSB (Saad *et al*., 2014). This initial decrease in mobility may contribute to repair using “local” donor sequences such as the sister chromatid, while increased local and global mobility following cell cycle arrest may facilitate interactions with alternative sequences that are less desirable templates but allow for DSB repair.

The outcome of homology search is also impacted by the regulation of strand invasion by the nucleoprotein filament as it samples potential templates. Factors such as the BLM helicase (Rqh1 in *S. pombe*) are thought to dissolve D-loops, thereby driving non-crossover repair events (Lorenz *et al*., 2014). Rqh1 likely promotes non-crossover products by favoring synthesis-dependent strand annealing (SDSA), in which strand invasion leads to new synthesis followed by dissolution of the D-loop, strand annealing that spans the initial DSB site, and repair (Symington *et al*., 2014; Symington, 2016). However, direct observation of this Rqh1 activity has not yet been possible.

Here, we describe the development of a microscopy-based assay in diploid fission yeast to determine the dynamics and kinetics of an engineered, interhomologue repair event. Although the initial distance between DSB and donor sequence predicts the time to their first physical encounter, it fails to predict the time to repair. Instead, repair efficiency is dictated by the number of strand invasion events, with most repair requiring multiple strand invasion cycles. In the absence of Rqh1, successful repair requires a single strand invasion event, suggesting that multiple strand invasion cycles reflect ongoing SDSA. This work therefore reveals the spatial and temporal events that influence homology-directed repair outcomes in living cells.

## Results and Discussion

### A microscopy assay to study interhomologue repair in living fission yeast

In order to monitor the timing and dynamics of homology search, we took advantage of a mating type mutant of *S. pombe* (*mat2-102;* Egel, 1973; Bodi *et al*., 1991) to generate stable diploids. In all cases, one of the haploid strains contains a site-specific HO endonuclease cut site adjacent to the *mmf1* gene, expresses Rad52(Rad22)-mCherry and has a floxed marker at the *urg1* gene that facilitates efficient Cre-mediated integration of the HO endonuclease such that it is regulated by the uracil-regulated *urg1* promoter (Watson *et al*., 2011). The other haploid strain has a 10.3 kb array of *lacO* repeats integrated adjacent to *mmf1* and expresses GFP-LacI (**Figure 1A**). Cells therefore have a single GFP focus and a diffuse distribution of Rad52-mCherry in the absence of HO endonuclease expression when visualized by fluorescence microscopy (Leland *et al*., 2018) (**Figure 1A**). We have shown previously in haploid cells that such a system induces a site-specific and irreparable DSB during S-phase on both replicated copies upon addition of uracil to the growth media (Leland *et al*., 2018). In this diploid system the induced DSB can undergo interhomologue repair (**Figure 1B**), with the DSB searching the nuclear volume and utilizing the homology near *mmf1* on the *lacO* array-containing homologous chromosome as the donor sequence (**Figure 1A**). As we observed previously in haploid cells (Leland *et al*., 2018), DSB induction and end resection lead to the recruitment of Rad52-mCherry, a proxy for the formation of the nucleoprotein filament that facilitates homology search and strand invasion, in ~15% of cells (**Supplementary Figure S1A**). While all cells display transient and dim Rad52-mCherry foci during S-phase (prior to cytokinesis), we hypothesized that the formation of a Rad52-mCherry focus at the site-specific DSB could be inferred by progressive and long-lived (>15 minutes) Rad52 loading induced at S phase. Indeed, cells without the induction of HO nuclease demonstrate only sporadic Rad52-mCherry loading (**Supplementary Figure S1B**). The percent of frames (taken every 5 minutes) in which a Rad52-mCherry focus is observed is significantly higher for cells (n=47) with HO nuclease induction than without (**Supplemental Figure S1C**). This interpretation was further validated experimentally (see below).

**Figure 1:**
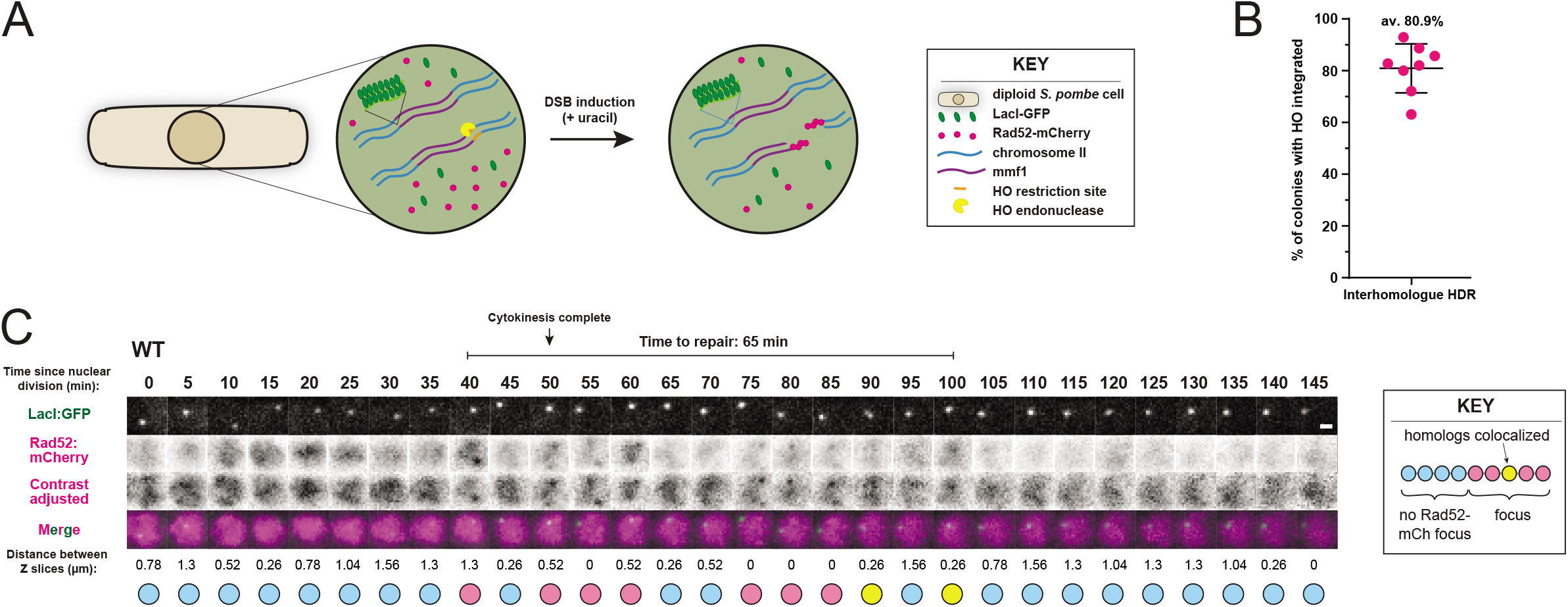
A fission yeast model system to monitor homology search during interhomologue repair in single, living cells. **(A)** Experimental design for the repair of a site-specific DSB in diploid fission yeast. A recognition site for the HO nuclease is integrated near the *mmf1* gene on one copy of Chr II. On the other copy of Chr II there is a *lac* operator array integrated ~ 5 kb from *mmf1*. The other assay components include expression of lacI-GFP, Rad52-mCherry, and HO nuclease from the uracil-regulated *urg1* promoter. **(B)** Interhomologue repair (mitotic recombination) is the dominant mode of homology-directed repair in diploid fission yeast. The proportion of cells expressing the HO endonuclease that undergo interhomologue repair, as determined by HO cut site marker loss assay (see Methods). Data from 8 biological replicates each containing between 50 and 200 colonies. **(C)** Efficient homology search and subsequent repair during interhomologue repair in fission yeast. Representative cell undergoing repair of the HO-induced DSB (see Supplementary Figure 2 for additional representative cells). Below the images the events are indicated as blue circles (no Rad52-mCherry focus), pink circles (Rad52-mCherry focus present but not colocalized with the donor), or yellow (Rad52-mCherry focus present and colocalized with the donor)(see Methods for details). Contrast of Rad52-mCherry signal adjusted according to the full histogram of intensities where indicated. Scale bar = 1μm.

An example of the time course of repair timing and chromatin dynamics within the 3D nuclear context is presented in **Figure 1C**. Images were acquired at 5 minute intervals for 3 hours after addition of uracil to induce expression of the HO nuclease. The lacI-GFP marking the donor sequence can be monitored throughout the movie. In this example, persistent Rad52-mCherry loading occurs at 40 minutes following nuclear division and persists up to 100 minutes following nuclear division (65 minutes total). Colocalization between the Rad52-mCherry loaded DSB and the donor sequence first occurs at 90 minutes post nuclear division, with Rad52 eviction 10 minutes later (100 minutes post nuclear division). The relationship between loss of a persistent Rad52-mCherry focus and repair was affirmed by monitoring subsequent cell division (see example, **Supplementary Figure S1D**).

As this system relies on inferring on-target, site-specific DSBs, we next carried out several controls to rigorously test if the dynamics we observe indeed reflect homology-directed repair and can be meaningfully interpreted. First, we determined the likelihood that the two *mmf1* loci would, at the diffraction limit of the light microscope, be found colocalized due to random fluctuations of the chromosomes in the absence of DSB induction. To this end, we generated a diploid strain in which a *lacO* array was integrated at both copies of *mmf1* (**Figure 2A, B**) and assessed the frequency at which the two *lacO* foci were found to be coincident. Under our imaging conditions, we find that the two *lacO*-GFP-LacI foci cannot be resolved in ~10% of frames during G2 (the cell cycle stage when we monitor repair (Leland *et al*., 2018), the majority of the *S. pombe* cell cycle) (**Figure 2C**). This is in stark contrast to the analysis of an aggregated cohort of WT cells with DSBs (n=21), in which Rad52-mCherry foci co-localized with the lacI-GFP-tagged donor sequence in ~35% of 5 minutes frames (**Figure 2C**). Thus, the majority of colocalization events between the Rad52-mCherry-loaded DSB and the donor sequence require the presence of the DSB.

**Figure 2:**
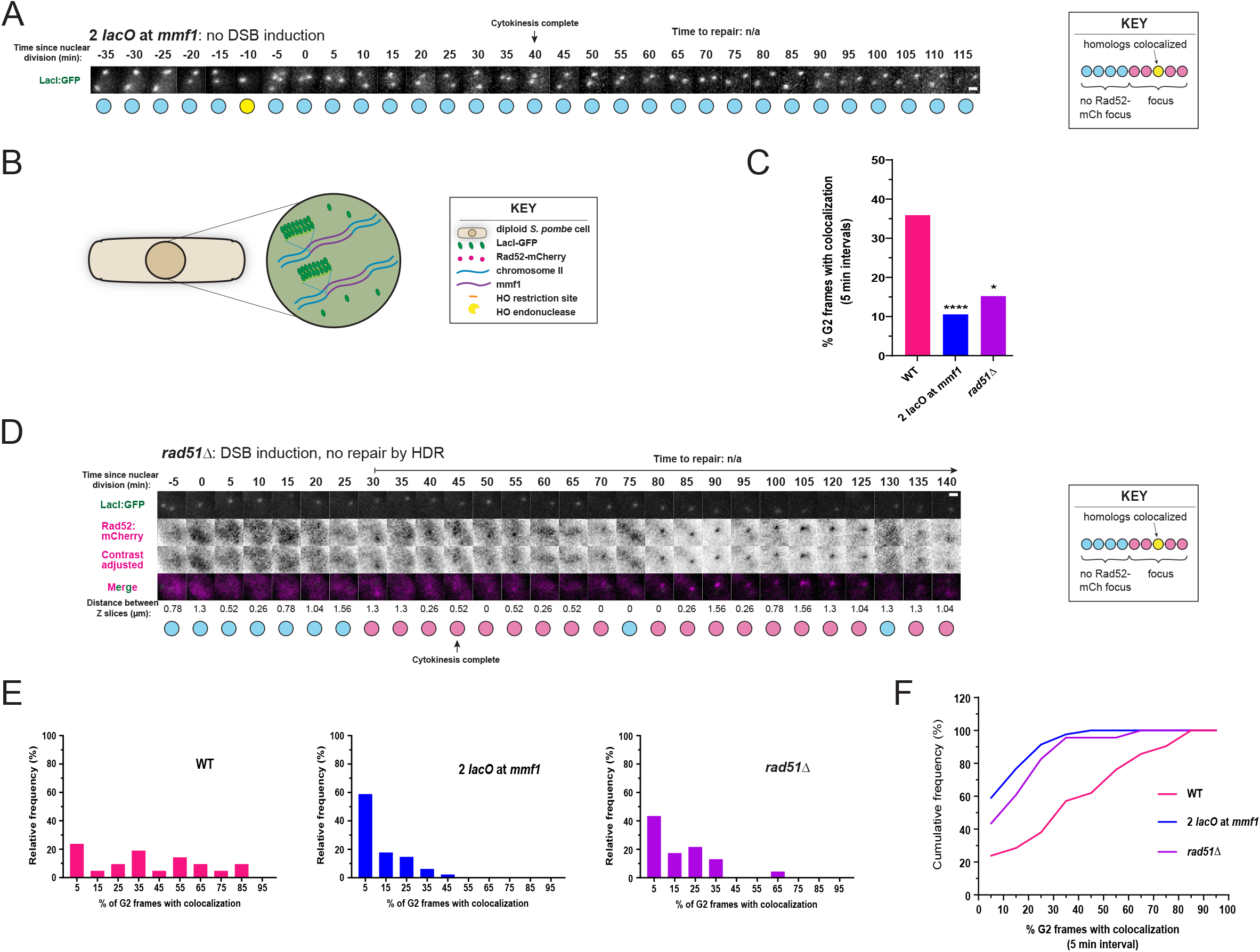
Colocalization of the DSB and donor sequence is driven by DSB formation and is Rad51-dependent. **(A)** ChrII homologs near the *mmf1* gene undergo minimal colocalization in the absence of an induced DSB. Z stack images of a nucleus from a representative 2 *lacO* at *mmf1* cell (see Methods and Figure 2B). Imaged as described in Methods, with 5 minutes between each time frame (columns) and labeled relative to nuclear division. Contrast adjusted to the full histogram of intensities where indicated. Scale bar = 1 μm. **(B)** Experimental design for monitoring of *mmf1* at both homologous chromosome II loci in the absence of DSB induction (2 *lacO* at *mmf1* background). On both copies of Chr II there is a *lac* operator array integrated ~ 5 kb from *mmf1* (see Methods), and lacI-GFP is expressed to visualize both homologs. The DSB was not induced with the HO/urg1 system. **(C)** Colocalization of homologs near *mmf1* is largely dependent on DSB induction and Rad51. Frames in which cells were in G2 phase were analyzed for colocalization of the DSB and donor (for WT and *rad51Δ*, only cells judged to have persistent, site-specific DSBs (see Methods) were included) and averaged as a total percentage across all cells. Colocalization is that of the DSB (Rad52-mCherry) and donor sequence (LacI-GFP bound to *lacO* repeats at *mmf1*) (WT and *rad51Δ*) or both chrII homologs in the absence of damage (2 *lacO* at *mmf1*). *p < 0.05, ****p < 0.0001, Kolmogorov-Smirnov test of cumulative distributions (of percentages from individual cells). WT: n=26, 2 *lacO* at *mmf1:* n=129, *rad51Δ:* n=23. **(D)** The DSB induced by HO endonuclease is persistent in *rad51Δ* cells. Z stack images of a representative *rad51Δ* induced cell imaged with 5 minutes between each time frame (columns) and labeled relative to nuclear division as described in Methods. Contrast of Rad52-mCherry signal adjusted according to the full histogram of intensities where indicated. Scale bar = 1μm. **(E-F)** Colocalization between the DSB and donor sequence is far more prevalent in WT cells than in *rad51Δ* cells or for cells with two *lacO* arrays at *mmf1* in the absence of damage. **(E)** Relative frequency histograms of percentages of G2 frames with colocalization in individual 2 *lacO* at *mmf1* control cells (n = 129), *rad51Δ* DSB cells (n = 23), and WT DSB cells (n = 21) (≥5 G2 frames per cell). Colocalization is for the DSB (Rad52-mCherry) and donor sequence (LacI-GFP bound to *lacO* repeats at *mmf1*) (WT and *rad51Δ*) or both chrII homologs in the absence of damage (2 *lacO* at *mmf1*). **(F)** Cumulative frequency histograms of data in Figure 2E.

To further test if the observed colocalization events are driven by strand invasion, we examined cells lacking Rad51, which is required for all homology search and strand invasion during HDR. Colocalization events between the induced DSB and donor sequence were strongly attenuated in *rad51Δ* cells (**Figure 2C and D**), nearly to the level observed in the absence of damage in the control 2 *lacO* cells (**Figure 2C**) despite persistent Rad52-mCherry loading. This suggests that most encounters between the DSB and donor sequence occur due to homology search via Rad51. We also analyzed the relative time of DSB-donor sequence colocalization in individual cells in all three conditions (**Figure 2E**). We observe that colocalization events in control 2 *lacO* cells without DNA damage and *rad51Δ* cells with DSB induction are short-lived compared to a broad distribution of lifetimes in WT cells, a conclusion reinforced by the difference in cumulative probability of colocalization frequency (**Figure 2F**).

### A site-specific DSB promotes multiple encounters with the homologous donor

The characteristic time required to successfully orchestrate HDR is fission yeast is still an outstanding question. Near complete recovery of a site-specific I-PpoI-induced DSB at a “generic” locus or at the rDNA repeats in fission yeast took place within 4 hours as measured by qPCR (Ohle *et al*., 2016), but no data was reported for times less than 4 hours. Other assay systems often employ donor sequences with only short-range homology to the DSB or utilize a mini-chromosome. By contrast, here we monitor repair between true homologous chromosomes. In addition, here we specifically monitor the time from the onset of long-range DSB end resection (~125 resected bps; Leland *et al*., 2018) to the eviction of Rad52, which corresponds more closely to the period of homology search. Consistent with this, we find that repair as defined here is highly efficient. The mean time to repair is ~50 minutes, although there is substantial cell-to-cell variation with a standard deviation of ~20 minutes (**Figure 3A**).

**Figure 3:**
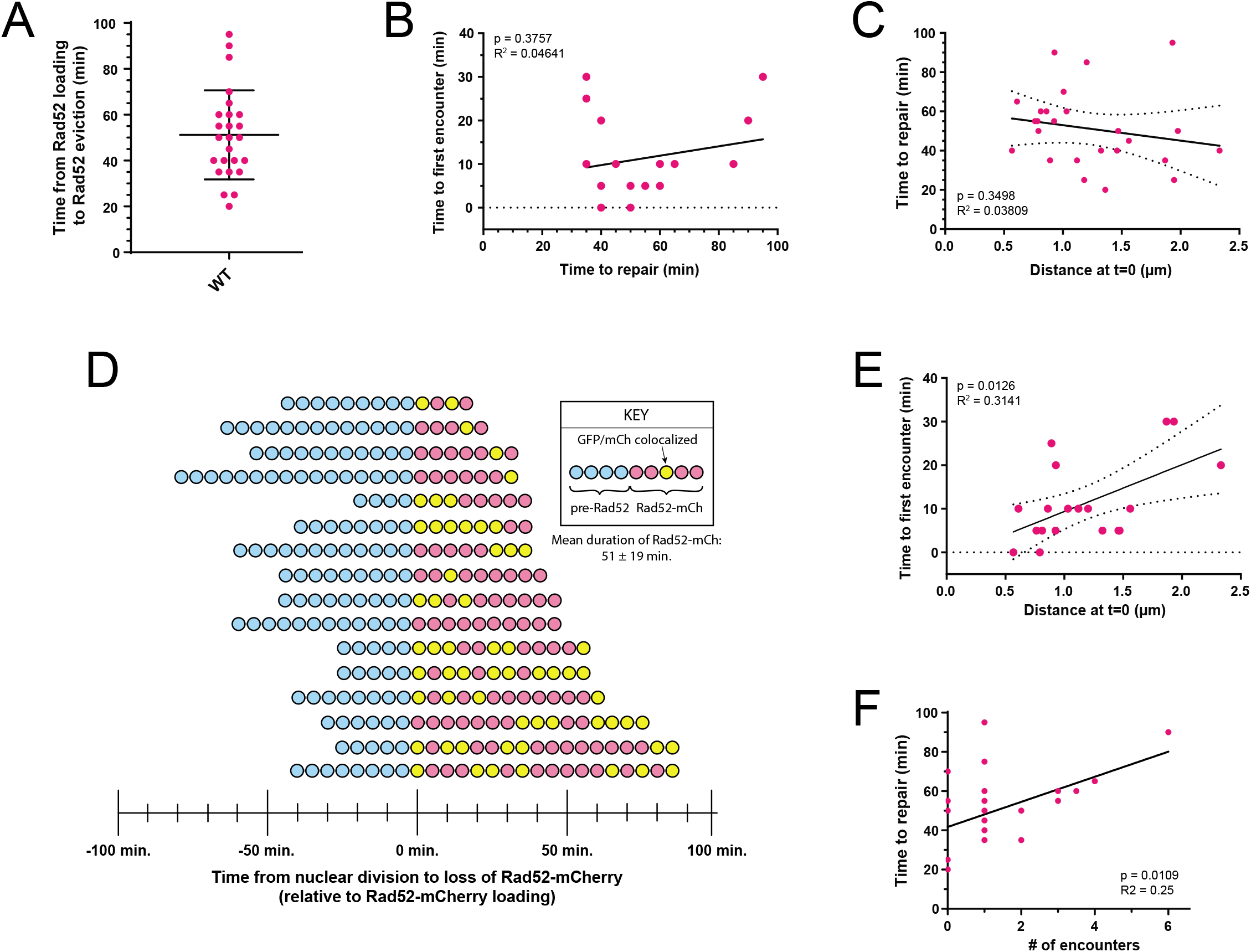
HDR in fission yeast frequently involves multiple encounters between the DSB and donor sequence. WT cells were imaged as described in Methods. Data points in **(A-C)** and **(E-F)** represent individual cells. **(A)** Repair of WT fission yeast cells is highly efficient in our induced DSB system. Time to repair was measured as the time in minutes from the first appearance of a site-specific DSB (persistent rad52-mCherry focus) to its disappearance for at least three consecutive frames (5 minute intervals, n = 25). Mean = 51.2, standard deviation = 19.4. **(B)** Timing of the first encounter between the DSB and donor sequence and timing of repair are not correlated. Time to first encounter is the difference between the first frame when Rad52-mCherry is visualized and the first colocalization event. Time to repair was measured as in Figure 3A. Linear regression: p value = 0.3757, R^2^ = 0.04641 (n = 16). **(C)** Timing of repair and initial distance are not correlated. Initial distance between the DSB and donor sequence was measured as the 3-D distance between the centers of the Rad52-mCherry (DSB) and lacI-GFP (donor) foci in the first frame after appearance of a site-specific (persistent) Rad52-mCherry focus. Time to repair was measured as in Figure 3A. Linear regression: p value = 0.3498, R^2^ = 0.03809 (n = 24). **(D)** Many WT DSB cells experience multiple colocalization events during repair, with variability in repair timing as well as number and length of colocalizations. Graph of colocalization events of Rad52-mCherry (DSB) and lacI-GFP (donor) foci in representative WT DSB cells. Each row represents one individual cell, and each circle represents a time point taken every 5 minutes. Blue circles: time from nuclear division to Rad52-mCherry loading. Pink circles: time from Rad52-mCherry loading to unloading for at least 3 consecutive frames. Yellow circles: Colocalization of the DSB (Rad52-mCherry) and donor (LacI-GFP bound to *lacO* repeats at *mmf1*) foci. See Supplementary Figure 2 for additional representative cells. **(E)** Timing of the first encounter between the DSB and donor sequence is correlated with their initial distance. Time to first encounter is the difference between the first frame when Rad52-mCherry is visualized and the first colocalization event. Initial distance was measured as in Figure 3C. Linear regression: p value = 0.0126, R^2^ = 0.3141 (n = 18). **(F)** The number of individual encounters is correlated with the timing of repair in individual cells. # of visualized encounters indicates the number of separate encounters (one or more consecutive frames (at 5 min. intervals) with colocalization) between the DSB and donor. Time to repair was measured as in Figure 3A. Linear regression: p value = 0.0109, R^2^ = 0.25 (n = 20).

Our initial expectation was that repair time corresponds to a single colocalization event reflecting strand invasion of the donor sequence by the DSB, new synthesis and ultimate repair. In this case, we would expect that (1) the time to the first encounter and the time to repair are correlated, if not equivalent, and (2) the initial distance between the loci and the time to repair are correlated(Lee *et al*., 2015). However, in this assay we observed that the time to the first encounter and the time to repair are not correlated (**Figure 3B**). Additionally, we found that the initial distance between the DSB and donor sequence does not correlate with repair time (**Figure 3C**), suggesting that an encounter *per se* is not the rate-limiting factor of homology search leading to repair. Instead, we frequently observe multiple colocalization events in individual cells over the course of DSB repair (**Figure 3D**). We therefore examined if the initial distance between the DSB and donor sequence correlates with the time to the initial colocalization event. Indeed, our analysis confirmed such a relationship (**Figure 3E**). Given that most colocalization events are Rad51-dependent (**Figure 2D-F**), we infer that many cells undergo multiple strand invasion events between the DSB and donor sequence prior to repair. If true, we would expect repair time to be tied to the number of strand invasion events. Indeed, we observe a positive correlation, supporting this interpretation (**Figure 3F**).

### Cells lacking Rqh1 display a bimodal repair phenotype and often repair after a single encounter between the DSB and donor

Based on the prevalence of multiple encounters between the DSB and donor sequence and variability in repair timing, we next considered if these kinetics reflect anti-recombination pathways that enforce HDR fidelity and/or non-crossover repair by SDSA. To address this, we tested the impact of deleting the *S. pombe* RecQ helicase, Rqh1, orthologous to human BLM. Rqh1 is established to dissolve D-loops (Van Brabant *et al*., 2000; Bachrati *et al*., 2006; Hope *et al*., 2007) and also contributes to DSB end resection in some contexts (Nanbu *et al*., 2015; Yan *et al*., 2019). However, we previously demonstrated that Rqh1 is dispensable for end resection in otherwise WT cells(Leland *et al*., 2018). Thus, the primary role(s) for Rqh1 in fission yeast HDR are downstream of resection and likely involve regulation of strand invasion structures.

In cells lacking Rqh1 we observe two distinct repair outcomes. In one subset of cells we observe very rapid repair (**Figure 4A**), while in another we observe highly persistent DSBs (**Figure 4B**). Indeed, the rate of productive repair within 90 minutes of initial Rad52-mCherry loading falls from over 65% in WT cells to ~40% in *rqh1Δ* cells (**Figure 4C**), suggesting that loss of Rqh1 negatively impacts repair as a whole. However, *rqh1Δ* cells that execute repair do so faster on average than for WT cells (**Figure 4D**). Given Rqh1’s role in D-loop disassembly, we next examined if the more rapid repair reflected a higher likelihood that a strand invasion event leads to repair. Indeed, we observe far fewer encounters between the DSB and donor in *rqh1Δ* cells that successfully repair, both in the population as a whole (**Figure 4E**) and within individual cells, where we often fail to visualize colocalization prior to repair within the 5 minute frame rate (**Figure 4F**). Thus, repair in *rqh1Δ* cells is bimodal, being either more efficient than in WT cells (often involving a single colocalization event) or failing entirely within our experimental observation window.

**Figure 4:**
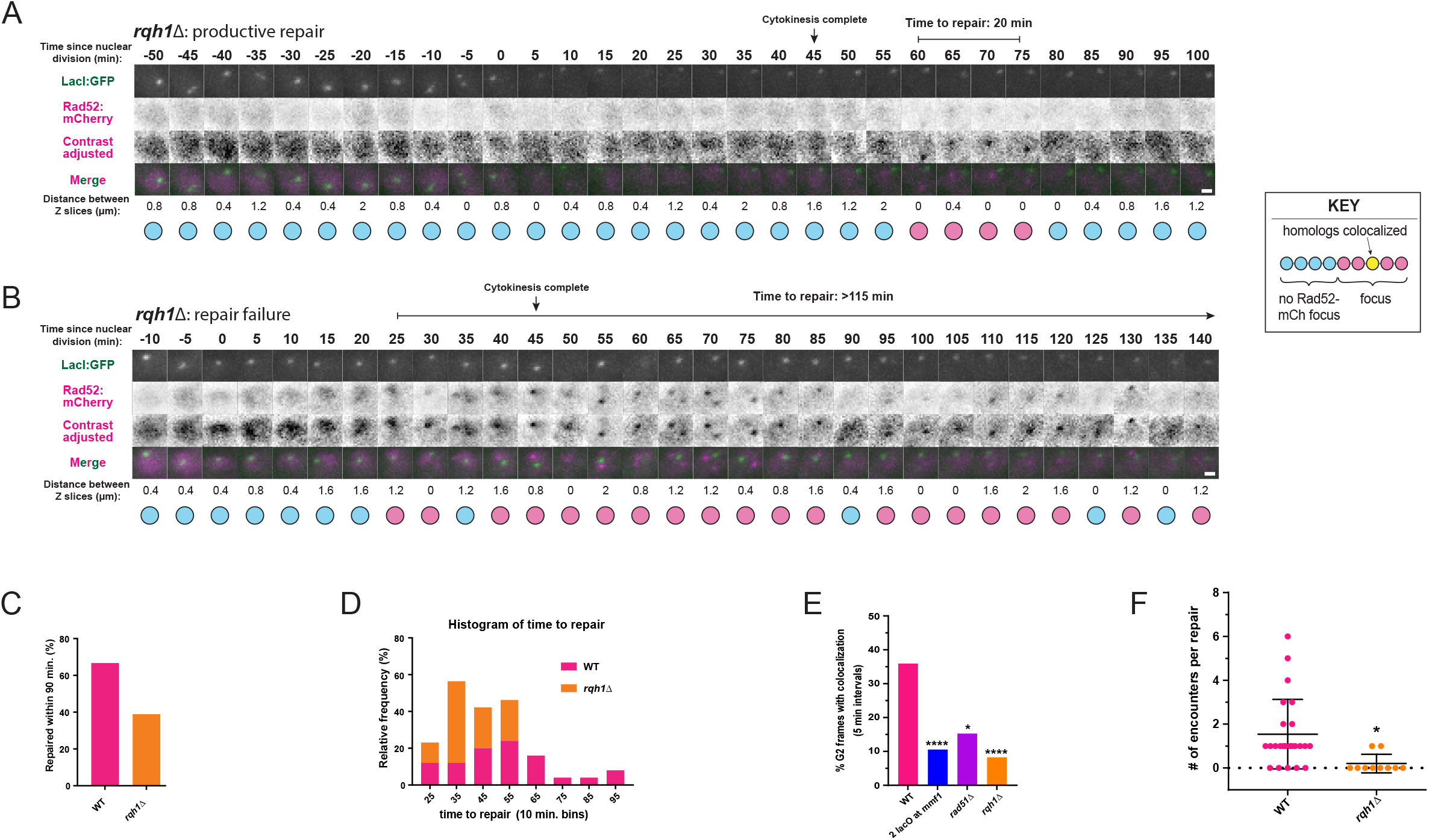
DSB cells lacking Rqh1 display a bimodal repair phenotype and have fewer encounters between the DSB and donor during repair. **(A)** Successful repair events are often relatively short in *rqh1Δ* DSB cells. Z stack images of a representative *rqh1Δ* DSB cell nucleus showing productive repair (persistent loss of Rad52-mCherry signal for at least 3 frames). Imaging as described in Methods with 5 minutes between each time frame (columns). Contrast of Rad52-mCherry signal adjusted to the full histogram of intensities where indicated. Scale bar = 1 μm. **(B)** Failure to repair DSBs efficiently is more prevalent in *rqh1Δ* cells with the induced DSB. Z stack images of a representative *rqh1Δ* DSB cell nucleus showing repair failure (persistence of Rad-52mCherry signal >90 min). Imaging as described in Methods\ with 5 minutes between each time frame (columns). Contrast of Rad52-mCherry signal adjusted according to full histogram of intensities where indicated. Scale bar = 1μm. **(C)** Cells lacking Rqh1 are less likely to undergo efficient DSB repair than WT cells. Total percentage of WT (n = 37) and *rqh1Δ* (n = 37) cells with an induced DSB that repair within 90 minutes. **(D)** Cells lacking Rqh1 have much shorter successful repair times. Stacked (no data hidden) relative frequency histograms of time to repair (10 minute bins) in WT cells (pink, n = 21; see Figure 3A) and *rqh1Δ* cells (orange, n = 16) with the induced DSB. **(E)** Cells lacking Rqh1 have a significantly smaller proportion of G2 frames with a colocalization per cell compared to WT. Quantification of colocalization of DSB (Rad52-mCherry) and donor sequence (LacI-GFP bound to *lacO* repeats at *mmf1*). Frames in which cells were in G2 phase were analyzed for colocalization of DSB and donor (for WT, *rad51Δ*, and *rqh1Δ*, only DSB cells were included) and assembled as a total percentage across all cells. Colocalization is that of the DSB (Rad52-mCherry) and donor sequence (LacI-GFP bound to *lacO* repeats at *mmf1*) (WT and *rad51Δ*) or both chrII homologs in the absence of damage (2 *lacO* at *mmf1*). *p < 0.05, ****p < 0.0001, Kolmogorov-Smirnov test of cumulative distributions (of percentages from individual cells). WT: n=26, 2 *lacO* at *mmf1:* n=129, *rad51Δ:* n=23, *rqh1Δ:* n=37. **(F)** Cells lacking Rqh1 have significantly fewer encounters between the DSB and donor per cell relative to WT. # of encounters per repair indicates the number of separate encounters (one or more consecutive frames at 5 min. intervals with colocalization) between the DSB and donor in individual WT (n = 25) or *rqh1Δ* (n = 10) cells. *p = 0.0143, Kolmogorov-Smirnov test of cumulative distributions.

## Conclusion

Taken together, our data indicate a highly efficient homology search in fission yeast. Surprisingly, we observe not one but multiple site-specific and Rad51-dependent colocalization events between the DSB and donor prior to successful repair. This suggests that 1) the first successful homology search event is not always followed by repair and/or 2) that multiple strand invasion events contribute to repair by synthesis dependent strand annealing. Notably, multiple encounters between a DSB and a homologous sequence has been proposed previously (Piazza *et al*., 2017; Piazza and Heyer, 2019) in the context of mating type switching in budding yeast (Houston and Broach, 2006). While we suggest that the observed dissolution of D-loops by Rqh1 likely reflects its contribution to promoting repair by SDSA, it may also facilitate rejection of strand invasion intermediates with non-homologous or homeologous sequences; the latter could explain why we often observe concomitant repair failure and lack of colocalization events in cells lacking Rqh1. Indeed, expression of mutated forms of Sgs1 (the ortholog of BLM and Rqh1) abrogated colocalization events between a DSB and the repair template in budding yeast (Piazza *et al*., 2017). More broadly, new insights into the highly transient nature of D-loop processing in budding yeast (Piazza *et al*., 2019) supports the possibility of short-lived encounters that are regulated by Rqh1. In patients with mutations in BLM, mitotic and meiotic crossover events are greatly increased, leading to genome instability and cancer predisposition among other symptoms (Arora *et al*., 2014). Our observations suggest that defects in the ability to promote non-crossover repair by SDSA combined with an accumulation of dead-end repair intermediates could both contribute to disease etiology, consistent with the observation that mutated alleles of Rqh1 lead to the “cut” phenotype in fission yeast treated with DNA damaging agents (Stewart *et al*., 1997).

Further study is also needed to fully define the relationship between genome organization and HDR efficiency and outcome. We find that the initial position of the DSB relative to the donor sequence had no bearing on overall repair duration, although DSBs that began closer to the donor sequence experienced a first colocalization more efficiently. Although a similar lack of correlation was recently described in a *trans* repair reporter assay for budding yeast NHEJ (Sunder and Wilson, 2019), studies of ectopic HDR have documented a correlation of initial position with repair efficiency in budding yeast (Agmon *et al*., 2013; Lee *et al*., 2015). While this could reflect inherent differences between model organisms, we also note that these studies leverage a relatively short homologous cassette inserted at ectopic sites rather than the homologous chromosome employed here. Moreover, our observations suggest that, although dependent on homology search, repair efficiency in this system is primarily dictated by the number of strand invasion events. One possibility is that while D-loop dissolution promotes SDSA, it also limits the extent of synthesis from a single homology search event. Thus, multiple stand invasion cycles may be necessary for the extent of synthesis required to span the initial DSB, thereby supporting subsequent strand annealing and repair.

---

**Supplementary Figure 1:**
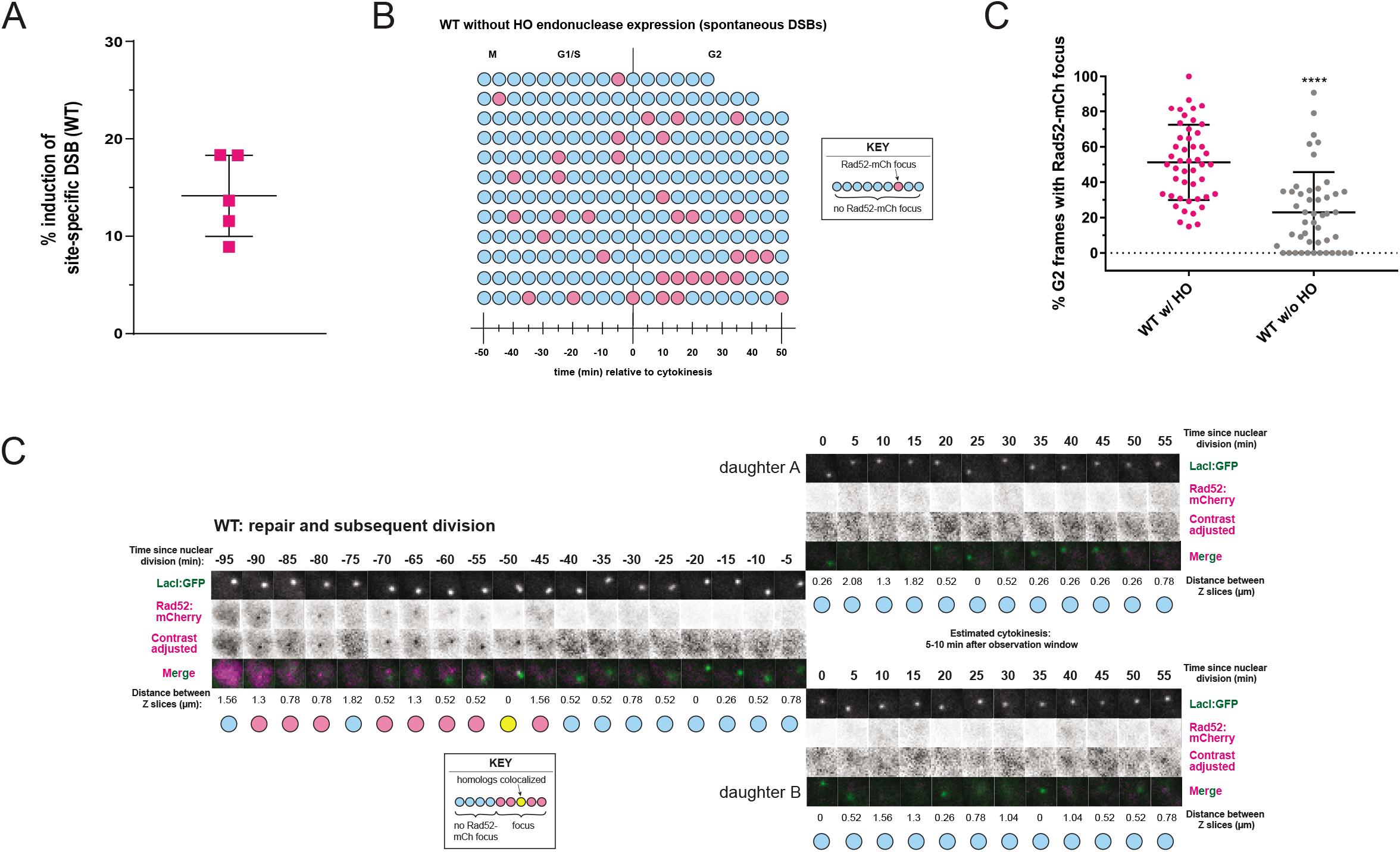
Induction of a site-specific DSB in WT cells during S phase has properties distinct from short-lived, non-specific DNA damage. **(A)** Proportion of WT cells with a site-specific DSB is similar to that seen previously (Leland *et al*., 2018) for the inducible HO/urg1 expression system. Proportions of WT cells with a sitespecific DSB from 5 technical replicates (4 separate inductions) (n > 200 per replicate except for a replicate of n = 101). **(B)** Spontaneous DSBs are short-lived and occur at random times in the cell cycle in WT cells without HO nuclease expression. WT cells were prepared and imaged as if induced but without transformation of the plasmid containing the HO endonuclease. Each row represents one individual representative cell, and each circle represents a time point taken every 5 minutes. Time points shown are between 50 minutes before and 50 minutes following cytokinesis, denoting the beginning of G2 (the observation window for the first two cells were shorter than 50 minutes following cytokinesis). Blue circles: nucleus did not contain a Rad52-mCherry focus in that frame. Pink circles: nucleus contained a Rad52-mCherry focus in that frame. **(C)** WT cells with HO-induced DSBs have a significantly greater proportion of G2 frames with a Rad52-mCherry focus than WT cells with spontaneous DSBs. Frames in which cells were in G2 phase were analyzed for the presence of a Rad52-mCherry focus (only cells with at least one frame with a Rad52 focus throughout the observation window were included). Data represent percentages from individual cells in G2 for at least 5 frames. ****p < 0.0001, Kolmogorov-Smirnov test of cumulative distributions (of percentages from individual cells). WT w/ HO: n=47, WT w/o HO: n=47. **(D)** Cells observed to induce the site-specific DSB followed by repair reenter the cell cycle as indicated by subsequent cell division, validating successful repair. Representative example of a WT cell in which cell division is observed after repair of the HO-induced DSB. Time before and after nuclear division is noted for each frame, and cytokinesis is estimated to have followed the observation window by 5 to 10 minutes. Images were acquired every 5 minutes (columns) (see Methods for details). Contrast of Rad52-mCherry signal adjusted according to the full histogram of intensities where indicated. Scale bar = 1 μm.

**Supplementary Figure 2:**
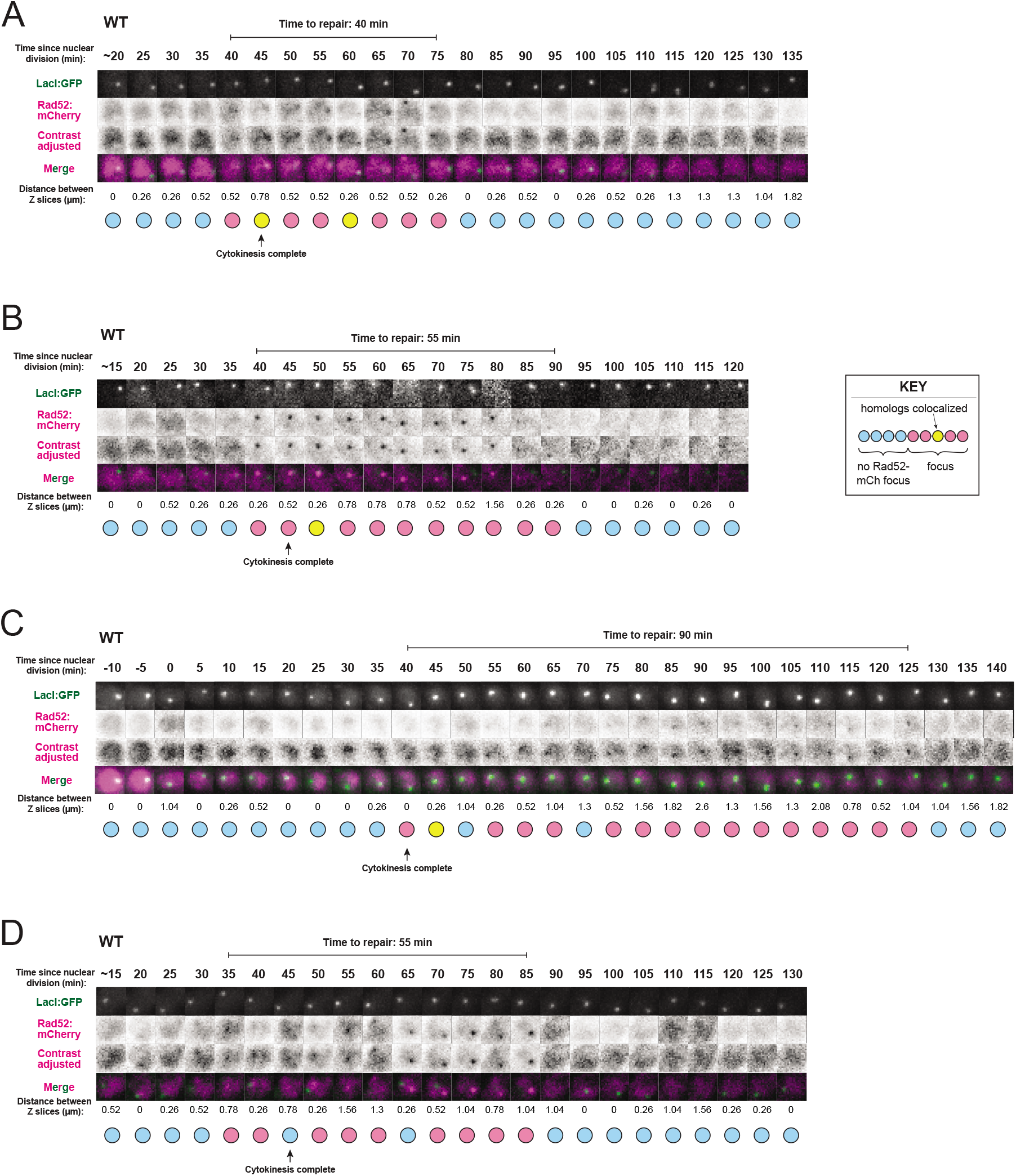
WT DSB cells exhibit a range of repair times, as well as number and length of colocalization events. **(A-D)** Representative cells undergoing repair of the HO-induced DSB. Repair times ranged widely in WT DSB cells (see **Figure 3A**). Additionally, cells might (**A**) (Figure 1C) exhibit multiple colocalizations during repair, (**B-C**) have single colocalizations with long periods of no colocalization, or (**D**) have no colocalization. Time of nuclear division was estimated based on cytokinesis in brightfield images (**A-B**), (**D**) or denoted as 0 minutes (**C**). Images were acquired every 5 minutes (columns) and are indicated relative to nuclear division (see Methods for details). Contrast of Rad52-mCherry signal adjusted according to the full histogram of intensities where indicated. Scale bar = 1 μm.

## Materials and Methods

### Yeast culture, strain construction and DSB induction

The strains used in this study are listed in Supplementary Table 1. *S. pombe* were grown, maintained, and crossed using standard procedures and media(Moreno *et al*., 1991). Gene replacements were made by gene replacement with various MX6-based drug resistance genes(Bähler *et al*., 1998; Hentges *et al*., 2005). In one haploid h-strain, the 10.3 kb LacO array was inserted between Mmf1 and Apl1 on the right arm of chromosome II (Chr II: 3,442,981) using a modified two-step integration procedure that first creates a site-specific DSB to increase targeting efficiency of linearized plasmid pSR10_ura4_10.3kb(Rohner *et al*., 2008; Leland *et al*., 2018). In another haploid *mat2-102* strain (competent to make a stable diploid when mated with an h-strain), a modified MX6-based hygromycin-resistance cassette containing the HO cut site was inserted between Apl1 and Mug178 on chromosome II (Chr II: 3,446,249). This insertion is 3.2 kb distal to the site of LacO insertion in the h-strain. DSB induction using the Purg1lox-HO system was performed as previously described(Leland and King, 2014; Leland *et al*., 2018).

### DSB induction using Purg1lox-HO

We used the uracil-responsive Purg1lox expression system, with slight modifications, to induce HO endonuclease expression and create site-specific DSBs at the HO cut site(Watt *et al*., 2008; Watson *et al*., 2011). We performed a fresh integration of the HO gene at the endogenous urg1 locus for each experiment in order to reduce long-term instability at the HO cut site or the development of HO resistance, presumably due to insertion/deletion events caused by basal expression levels of HO. The pAW8E*Nde*I-HO plasmid (a gift from Tony Carr) was transformed into *S. pombe*, which were then plated onto EMM-leu+thi-ura plates (-leucine: plasmid selection; +thiamine: Pnmt1-Cre repression; -uracil: Purg1lox-HO repression). After 4–6 days of growth at 30°C, 20-60 individual colonies were combined to obtain a reproducible plasmid copy number across the population. Cre-mediated HO gene exchange at the endogenous Urg1 locus (urg1::RMCEkanMX6) was induced by overnight culture in EMM-thi-ura+ade+NPG media (thiamine: expression of Cre from pAW8ENdeI-HO; -uracil: Purg1lox-HO repression; +0.25 mg/mL adenine: reduce autofluorescence; +0.1 mM n-Propyl Gallate (NPG): reduce photobleaching in microscopy experiments, prepared fresh). The following day, site-specific DSBs were induced in log-phase cultures by the addition of 0.50 mg/mL uracil. This induction strategy resulted in ~15% of cells making a DSB within ~2 hr (Supplementary Figure 1A).

### Microscopy

All images were acquired on a DeltaVision widefield microscope (Applied Precision/GE) using a 1.2 NA 100x objective (Olympus), solid-state illumination, and an Evolve 512 EMCCD camera (Photometrics). Slides were prepared ~10-20 min after adding 0.50 mg/ml uracil to log-phase cultures to induce HO endonuclease expression and DSB formation. Cells were mounted on 1.2% agar pads (EMM +0.50 mg/mL uracil, +2.5 mg/ml adenine, +0.1 mM freshly prepared NPG) and sealed with VALAP (1:1:1 vaseline:lanolin:paraffin). Image acquisition began between 40 and 80 min after uracil addition. Imaging parameters for all microscopy assay data acquisition were as follows. Transmitted light: 35% transmittance, 0.015 s exposure; mCherry: 32% power, 0.08 s exposure; GFP: 10% power, 0.05 s exposure. At each time point (every 5 min for 2.5-4 hr), 25 Z-sections were acquired at 0.26mm spacing (16 Z-sections were acquired at 0.42mm spacing to mitigate photobleaching in some samples).

### Image analysis

For the microscopy assay of interhomologue repair, data were acquired for each cell cycle individually, including time of nuclear division, time of cytokinesis, frames in which Rad52-mCherry focus was visible and frames in which Rad52-mCherry focus colocalized with the LacI-GFP focus at the diffraction limit (in the case of the 2 *lacO* at *mmf1* strain (**Figure 2A-B**), colocalizations between both LacI-GFP foci were recorded instead). Time to repair was measured as the time in minutes from the first appearance of a site-specific DSB (persistent rad52-mCherry focus) to its disappearance for at least three consecutive frames. Only sitespecific DSBs (defined as Rad52-mCherry focus persistence for at least 4 frames that began in late S or early G2 phase) were considered, since spontaneous DSB events can occur within the genome especially in G1 and early S phase (see **Supplemental Fig. 1B-C**). Fields were analyzed manually, using the same contrast settings for mCherry and GFP channels for consistency. Images from representative cells for each strain (**Figure 1C, 2A, 2D, 4A-B, Supplementary Figure 1D, Supplementary Figure 2A-D**) were prepared using ImageJ macros to automate merge and montage image creation using the same gate size (height and width), while allowing for manual selection of the Z plane and centering on the nucleus. For visual clarity, the contrast of some images was adjusted according to the histogram using Levels sampling functions of Adobe Photoshop (2018) to set the darkest pixel as black and the brightest pixel as white. Merged images are either max projection or single planes with Rad52-mCherry in focus for visual clarity. Distance between Z slices for each frame is the distance in Z between the Z slice containing the center of the LacI-GFP focus and the Z slice containing the center of the Rad52-mCherry focus (or the center of the nucleus (denoted by middle Z slice of diffuse Rad52-mCherry signal) in frames with no Rad52-mCherry focus).

Data were plotted and analyzed using GraphPad Prism 7.01. Percentages of G2 frames with colocalization from individual cells were analyzed using the Kolmogorov-Smirnov test of cumulative distributions (**Figure 2C and 4E**, p value denoted by asterisks and average plotted), relative frequency histograms (**Figure 2E**) and cumulative frequency histograms (**Figure 2F**). Linear regressions (**Figure 3B-C, E-F**) were calculated using default Prism settings. Dotted lines (**Figure 3C and E**) represent 95% confidence intervals. The number of encounters per repair (**Figure 4F**) were analyzed using the Kolmogorov-Smirnov test of cumulative distributions (mean and standard deviation plotted).

### Marker loss assay

To examine repair outcome of the DSB in our system results based on sequence changes resulting from different repair pathways at the HO cut site, we performed a marker loss assay to assess the proportion of induced cells in which the MX6-based drug resistance gene(Bähler *et al*., 1998; Hentges *et al*., 2005) was lost due to use of the donor sequence during HDR. DSB induction was performed on WT diploid *S. pombe* cells as described above. At 2 hours following induction in log phase (growth for 2 hours in EMM-ura+ade+NPG with uracil added), cells were resuspended in EMM-ura media and plated to YE5S at 1:1000 (n=3), 1:2000 (n=3) and 1:5000 (n=2) dilutions. After 24 hours, YE5S plates were replica plated to YE5S+Kanomycin (at HO cut site - lost when the DSB is repaired using the homologous donor) and YE5S+Hygromycin (at *urg1::RMCE* - lost when the pAW8E*Mde*I-HO plasmid is flipped in via Cre recombination prior to induction). Colonies were counted with a Bio-Rad Molecular Imager VersaDoc (total colony count between 50 and ~160 cells per YE5S plate). Percentage of cells from each YE5S plate that had repaired by interhomologue HDR was calculated as (%Kan sensitive colonies/%Hyg sensitive colonies)*100. Data along with mean and standard deviation were plotted using GraphPad Prism 7.01.

**Supplementary Table 1:** Strains used in this study

| Strain | Description | Complete Genotype |
| --- | --- | --- |
| MKSP2230 | WT diploid with lacO repeats on both ChrII copies (control: no HO integration/DSB formation) | *mat2-102/h- leu1-32/leu1-32 ura4-D18/ura4-D18 his7+:P<sub>Dis1</sub>-GFP-LacI-NLS/ his7+:P<sub>Dis1</sub>-GFP-LacI-NLS ChrII:3442981::Ura4-10.3kbLacO/ ChrII:3442981::Ura4-10.3kbLacO ChrII:3446249::HOcs-hphMX6/WT Rad52-mCherry::natMX6/Rad52-mCherry::natMX6 urg1::RMCE-kanMX6/URG1* |
| MKSP2450 | WT homology search assay (mated with MKSP2489 to make diploid) | *mat2-102 leu1-32 ura4-D18 Rad52-mCherry::natMX6 ChrII:3446249::HOcs-kanMX6* |
| MKSP2489 | WT homology search assay (mated with MKSP2450 to make diploid) | *h- leu1-32 ura4-D18 his7+:P<sub>Dis1</sub>-GFP-LacI-NLS ChrII:3442981::Ura4-10.3kbLacO urg1::RMCE-kanMX6* |
| MKSP2632 | *rqh1Δ* homology search assay (mated with MKSP2788 to make diploid) | *h- leu1-32 ura4-D18 his7+:P<sub>Dis1</sub>-GFP-LacI-NLS ChrII:3442981::Ura4-10.3kbLacO urg1::RMCE-kanMX6 rqh1::hphMX6* |
| MKSP2709 | *rad51Δ* homology search assay (mated with MKSP2736 to make diploid) | *h- leu1-32 ura4-D18 his7+:P<sub>Dis1</sub>-GFP-LacI-NLS ChrII:3442981::Ura4-10.3kbLacO urg1::RMCE-kanMX6 rad51::hphMX6* |
| MKSP2736 | *rad51Δ* homology search assay (mated with MKSP2709 to make diploid) | *mat2-102 leu1-32 ura4-D18 Rad52-mCherry::natMX6 ChrII:3446249::HOcs-kanMX6 rad51::hphMX6* |
| MKSP2788 | *rqh1*Δ<br>homology<br>search assay<br>(mated with<br>MKSP2632 to<br>make diploid) | *mat2-102 leu1-32 ura4-D18 Rad52-*<br>*mCherry::natMX6 ChrII:3446249::HOcs-kanMX6*<br>*rqh1::hphMX6* |
| MKSP3038 | WT marker loss<br>assay (mated<br>with MKSP2450<br>to make diploid) | *h- leu1-32 ura4-D18 his7+:P<sub>Dis1</sub>-GFP-LacI-NLS*<br>*ChrII:3442981::Ura4-10.3kbLacO urg1::RMCE-*<br>*hygMX6* |

